# Deep learning based decoding of local field potential events

**DOI:** 10.1101/2022.10.14.512209

**Authors:** Achim Schilling, Richard Gerum, Claudia Boehm, Jwan Rasheed, Claus Metzner, Andreas Maier, Caroline Reindl, Hajo Hamer, Patrick Krauss

## Abstract

How is information processed in the cerebral cortex? To answer this question a lot of effort has been undertaken to create novel and to further develop existing neuroimaging techniques. Thus, a high spatial resolution of fMRI devices was the key to exactly localize cognitive processes. Furthermore, an increase in time-resolution and number of recording channels of electro-physiological setups has opened the door to investigate the exact timing of neural activity. However, in most cases the recorded signal is averaged over many (stimulus) repetitions, which erases the fine-structure of the neural signal. Here, we show that an unsupervised machine learning approach can be used to extract meaningful information from electro-physiological recordings on a single-trial base. We use an auto-encoder network to reduce the dimensions of single local field potential (LFP) events to create interpretable clusters of different neural activity patterns. Strikingly, certain LFP shapes correspond to latency differences in different recording channels. Hence, LFP shapes can be used to determine the direction of information flux in the cerebral cortex. Furthermore, after clustering, we decoded the cluster centroids to reverse-engineer the underlying prototypical LFP event shapes. To evaluate our approach, we applied it to both neural extra-cellular recordings in rodents, and intra-cranial EEG recordings in humans. Finally, we find that single channel LFP event shapes during spontaneous activity sample from the realm of possible stimulus evoked event shapes. A finding which so far has only been demonstrated for multi-channel population coding.

## Introduction

How is sensory input and information processed in the cerebral cortex? To answer this question, a lot of effort has been undertaken to measure brain activity with growing temporal and spatial resolution. Indeed, nowadays due to the enormous progress of fMRI, we know very exactly were certain processes take place. Thus, it is possible to scan the whole brain on a millimeter-scale within seconds [1].

On the other hand, to understand what is exactly happening in the brain when sensory stimuli are processed, an in-depth analysis of temporal cues of neural activity in the brain is necessary. However, it is still not fully clear how to take advantage of the growing temporal resolution of modern neuro-imaging techniques such as MEG/EEG, and intra-cranial multi-unit recordings. Thus, surface based methods like EEG and MEG provide a poor spatial resolution and signal-to-noise ratio, as the source of the signal and the recording site are divided by the skull. The surface electrodes, or SQUIDs respectively, are used to record the sum-activity of millions of neurons [2, 3]. Furthermore, the poor signal-to-noise ratio results in the issue that meaningful data can only be extracted, by averaging over many trials for evoked activity (for auditory evoked potentials ≥ 100, see [4, 5]), or by extracting spectral cues through Fourier-filters [6, 7]. However, as Waterstraat and co-workers state the brain is a *”single trial processor”*, that has been shaped by evolutionary processes [8].

The only way to increase the signal-to-noise ratio in order to extract meaningful data from single-trials are intra-cranial recordings. However, these experiments are limited to animal studies as the intra-cranial recording is an invasive method. However, in epilepsy diagnostics sometimes intra-cranial electrodes are implanted in human patients’ brains in order to exactly localize the epilepsy focus (iEEG, [9]). Usually, these rare recordings are additionally used to tackle scientific questions [10–12]. In contrast, to surface recordings on top of the skull such as EEG or MEG, LFP events from intra-cranial recordings have a far better resolution since they are mainly produced by three orders of magnitude smaller neuron ensembles [13]. Thus, the radius, within which neurons significantly contribute to the LFP is only in the range of a few hundreds of micrometers [14, 15]. Furthermore, LFP shapes contain information on the proportions of the contributing sources of extra-cellular currents as well as on the conductive properties of the surrounding brain tissue [11]. Thus, the low frequency parts of the LFPs are induced by synaptic activity, inducing trans-membrane currents, which lead to electrical dipoles. These dipoles can be measured using intra-cranial electrodes [11]. Although fast action potentials contribute to the high-frequency components of LFPs, the main contribution comes from dendritic currents [16–18]. Therefore, LFPs are typically regarded as a measure of the input signal to the neurons to be measured [17]. It has to be stated that further effects such as calcium spiking, gap junctions, and neuron oscillations play a role for LFP events [11].

Since the spectral cues as well as the shape of LFP events contain a lot of information on geometry, properties and activity of the neuron ensembles, the investigation of LFPs in the sense of sensory processing and cognition is a promising approach. Thus, LFPs are for example used to implement brain computer interfaces for locked-in patients [19].

Despite the fact that single trial LFP events are a valid measure for local neural population analysis [20], studies on single-trial LFP events analysis are rare. Thus, a better understanding of LFP events could e.g. help to make major progress in understanding cognitive processes during sleep [21]. Unfortunately, in contrast to multi-unit activity (MUA), the shapes of LFP events are more versatile. Hence, it is far more difficult to automatically cluster, respectively classify, the different LFP shapes compared to spiking activity (for spike clustering see e.g. [22]). However, it has already been demonstrated that wavelet analysis could be used to extract meaningful features from single trial LFP events [23]. Furthermore, an algorithm to automatically detect and calculate three pre-defined features (latency, amplitude, and rebound) of single trial LFP events has been developed [24].

Nevertheless so far, more advanced evaluation techniques such as machine learning based pattern recognition, especially using deep neural networks, has been applied only rarely to detect and characterize single trial LFP events. Thus, more coarse-grained features such as spectral cues have been analyzed using machine learning [25]. Nurse and co-workers used a top-down approach to classify LFP data by training a convolutional neural network on raw data [26]. They argue that the bottleneck in processing LFPs using machine learning is to find the right hand-crafted features [26].

However, such top-down approaches like e.g. training classification networks on raw LFP data, respectively events, have also major drawbacks compared to parameter-free and data-driven bottom-up approaches [27]. Although, classifyer approaches approach are promising to develop e.g. brain computer interfaces, the neuroscientific value of these networks is poor: Whereas we can derive statements on certain features of the LFP signal regarding our pre-defined labels, all the other information, which is not needed for classification according to the pre-defined labels is lost.

Therefore, a valid approach to deal with high-dimensional electrophysiological data sets, is to apply unsupervised machine learning algorithms. These algorithms are used to extract recurring patterns from the signal and to reduce the dimensionality of the signal for the purpose of visualizations, being better interpretable for humans [28–30]. Indeed, recently some studies have been published, which applied unsupervised machine learning methods on electro-physiological data. For instance, Mackevicius and coworkers developed an unsupervised training algorithm to extract non-redundant sequences of the neurophysiological data [31]. In addition, Hardcastle and coworkers [32] applied an auto-encoder network on peripheral nervous system recordings (for further example see also [33]). Note that, some scientific approaches used the unsupervised trained networks the other way around, namely as models for the nervous system (see e.g. [34]).

Here, we use an auto-encoder network to cluster different LFP shapes. Therefore, we extract the LFP events by searching for local minima in the signal stream and applying advanced thresholding and filtering techniques. The extracted LFP events are encoded to identify certain clusters representing different LFP shapes. For the resulting low-dimensional representations the term embeddings has been coined [35–37]. To increase interpretability, we decode the embeddings after we have identified different clusters, thereby reverse-engineering the prototypical LFP shape of each cluster. Furthermore, we provide evidence that the LFP shape may be used to distinguish between different stimulus frequencies for evoked activity and is a highly reproducible marker for the direction of information flow within the cerebral cortex for both evoked and spontaneous activity. In addition, we demonstrate that the maximum of the cross-correlation is linked to the shape of the LFP events. Finally, we show that single channel LFP event shapes during spontaneous activity sample from the realm of possible stimulus evoked event shapes. A finding which so far has only been demonstrated for multi-channel population coding. To put it in a nutshell, the self-supervised auto-encoding approach can be used to extract valuable features from intra-cranial recordings in rodents and from iEEG data recorded in human epilepsy patients.

## Methods

### Rodent Data Acquisition

#### Surgery

For the craniotomy, the Mongolian gerbils were put under deep Ketamin-Xylazine anesthesia and kept on a controlled heating pad to guarantee a constant body temperature. After hair removal using an electric razor the scalp was removed using sharp micro-scissor. The skull is cleaned with a drill and tweezers. Four screws (M2, 2 mm) are implanted frontally and caudally from bregma, which serve as base for the head fixation. As head-fixation a small elbow is glued with dental cement to the skull and the fixation screws. A rectangular trepanation of approximately 4 mm × 4 mm is opened located between ear and the eye contra-laterally to the ear to be measured. The dura is removed using a fine needle and micro-tweezers. The trepanation is covered with NaCl solution to prevent the brain from drying out.

#### Extracellular Recordings

After the craniotomy the animal is placed in an anechoic chamber on a controlled heating pad and the head is fixed by screwing the vertical part of the elbow to a aluminium rod. The distance of the loudspeaker and the ear of the animal is X cm. For extracellular recording we use 16 electrode arrays (Clunbury Scientific (Bloomfield Hills), 0.5 MΩ, layout:4 × 4, spacing; 500 *μ*m). The electrode is inserted in the auditory cortex of the animal using a manual micro-cotroller. The insertion depth of the electrode array is between 500 *μ*m and 1 mm to record signals from the granular and infra-granular layers, where we expect significant spiking activity and high field potential amplitudes. The neural signal is recorded using the CerePlex system from Blackrock Neurotech (Salt Lake City, USA). The neural signal is digitized directly at the recording site with the Cereplex *μ* headstage. We recorded all signals using the maximum sampling rate of 30 kHz. To divide between LFP and spike signal during recording, for LFPs a 250 Hz low-pass filter was applied, whereas spiking activity was assumed to occur in the range between 250 Hz and 2.5 kHz. The spikes were sorted online by manually set thresholds.

#### Auditory Stimulation

For auditory stimulation we used a 3 way loudspeaker (MAC, Racer 320). We calibrated the loudspeaker by measuring the output loudness for frequencies between 500 Hz and 19 kHz (half octave steps). For each frequency a correction factor was calculated. The loudspeaker is driven by a 24 bit external sound card (ASUS Xonar MKII 7.1) connected to the computer via USB. The soundcard output is amplified using an audio amplifier (Amp75). To ensure exact alignment of the auditory stimuli with the recorded neural signal, the soundcard is also used as trigger pulse generator. Thus, a second soundcard channel sends trigger pulses to an analog input channel of the recording setup (cf. also [35, 38]). To analyze the tuning of the neurons, we presented 50 ms pure tone stimuli with inter-stimulus intervals of 300 ms. Click sounds were prevented by adding sine^2^-ramps of 5 ms duration to the pure tone stimuli. The presented stimulus frequencies lie in the range of 500 Hz and 20 kHz (half octave steps). Stimulus intensities and frequencies were presented pseudo-randomly. Depending on the hearing ability of the animals, 5 different sound pressure levels were presented (10 dB steps) starting from 110 dB SPL (90 dB, 70 dB respectively).

### Human Data Acquisition

The intracranial EEG (iEEG as EEG recorded via stereotactically implanted depth electrodes, SEEG) originated from a patient suffering from epilepsy, who underwent presurgical epilepsy evaluation due to a drug refractory focal epilepsy. Depths electrodes were implanted in the auditory cortex of the right temporal lobe solely because of clinical reasons. In this study, we show the data of 4 platin electrodes (RTB-1-4) with an impedance lower than 10 kΩ, which were measured against a reference electrode placed on top of the scalp (CPz). Due to the fact that there are not many patients, who receive electrodes in the auditory cortex we are data limited.

### Data Analysis

#### Computational Resources

The complete software used for the project was written in Python 3.8 using the NumPy library ([39]). The graphical user interface was designed and implemented using PyQt5 [40]. The neuronal recordings were imported and converted using the ’brpylib’-library provided by Blackrock Neurotech [41]. Basic data processing and filtering was done using the SciPy-library and especially the ’signal’-module [42]. Machine learning was implemented in Keras [43] with the TensorFlow backend [44]. All simulations and evaluations were performed on a stadard desktop computer. Plotting was done using the Matplotlib library [45] in combination with the pylustrator add-on [46].

#### Neural Networks

The used neural networks in this study were all implemented using Keras with the tensorflow backend [43, 44]. The auto-encoder was trained on single channel LFP events from a 1 h spontaneous activity recording of only one animal. Thus, first local minimum technique (descibed above) was used to find the LFP events. These LFP events were then used to train the auto-encoder. Test-data from other recordings and other animals were used to prove the validity of the training. Before used for training the LFP-event were down-sampled to 100 Hz and cut into 500 ms junks (50 values). The 50-dimensional input vector is expanded by a factor of 14 to 700 dimensions in the first hidden layer of the network. It has be shown that dimensionality expansion can lead to a huge benefit for data processing in neural networks [47, 48]. The bottleneck layer has only 3 dimensions. This number was chosen as it is the highest dimensionality, which can be easily visualized (for details see Tab. 1). The batch-size for training was 500 and the training iterations were 1000. During training the data was randomly shuffled.

**Table 1:** Auto-Encoder Network (epochs = 1000, batch-size = 500)

| Layer | Activation | Output Shapes | Parameters |
| --- | --- | --- | --- |
| Input Layer | ReLu | (None, 50) | 0 |
| Dense | ReLu | (None, 700) | 35700 |
| Dropout (0.3) |  | (None, 700) | 0 |
| Dense | ReLu | (None, 3) | 2103 |
| Dense | ReLu | (None, 700) | 2800 |
| Dropout (0.3) |  | (None, 700) | 0 |
| Dense | ReLu | (None, 50) | 35050 |

In order to measure the loss of meaningful information caused by the auto-encoding procedure, we trained an additional feed-forward network. In particular, we used a 1D convolutional neural network consisting of two convolutional layers, one max-pooling layer, one fully connected (dense) layer, and finally a softmax layer for classification (for details see Tab. 2). This network was trained on the classification of LFP events according to 4 different stimulus frequencies, hence the output vector is 4-dimensional. The number of iteration was set to 10,000, but for the classifier the early stopping technique was applied.

**Table 2:** Classifier Network

| Layer | Activation | Output Shapes | Parameters |
| --- | --- | --- | --- |
| Input Layer |  | (None, 50, 1) | 0 |
| Convolution 1D | ReLu | (None, 41, 60) | 330 |
| Dropout (0.3) |  | (None, 41, 60) | 0 |
| Convolution 1D | ReLu | (None, 32, 30) | 18030 |
| Max Pooling 1D |  | (None, 1, 30) | 0 |
| Flatten |  | (None, 30) | 0 |
| Dropout (0.3) |  | (None, 30) | 0 |
| Dense | ReLu | (None, 20) | 620 |
| Dense | Softmax | (None, 4) | 84 |

#### Cross-Correlation Analysis

For correlation analysis we used the normalized cross correlation [49]. Thus, the LFP event of the reference channel and the LFP-events of the neighbouring channels to be tested were z-scored individually (subtract mean and divide by standard-deviation). After that, the cross-correlation was calculated using the correlate function from numpy [39]. The sum is then divided by *n_o_* representing the number of summands. Thus, this definition results in a normed cross-correlation and an auto-correlation of one for no lag time (time delay) *τ* = 0.

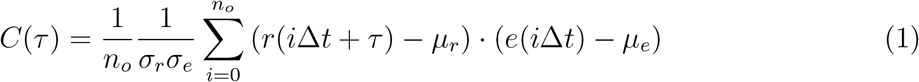

(*C*(*τ*): zero-normed cross correlation, *n_o_*: number of summands which is the number of overlapping samples, *r*(*i*∆*t*): reference channel value at time point *i*∆*t*, *i* ∈ {−*n_o_, n_o_* + 1…, *n_o_* − 1, *n_o_*}, ∆*t* =1 ms, *μ_r_*: mean of all values only for one LFP event in reference channel, *τ* : time delay, *τ* ∈ [−50 ms, 50 ms], *e*(*i*∆*t* +*τ*): value of tested channel at time point *e*(*i*∆*t* +*τ*), *μ_e_*: mean of test channel)

## Results

In the following paragraphs, we describe how to identify and group LFP events in a continuous LFP data stream.

### Detection of LFP events

In standard evaluation pipelines and especially in settings, where sensory stimuli are presented, LFP responses are averaged over several trials to remove the measurement noise. However, averaging over trials has several drawbacks such as the effect that inter-trial differences vanish and that always a well-defined stimulus and a time trigger is needed to group and align the recorded data. However, when the signal-to-noise ratio is large enough, then it is not necessary to average over several trials in intra-cranial recordings as the single trail events are already good enough to extract meaningful information. Thus, as shown in Fig. 1a, single pure-tone stimuli of varying frequency and loudness do already evoke clear LFP responses. In most studies, the LFP responses as well as the spike rates are averaged to calculate spectro-receptive fields (STRFs, Fig 1b) for different stimulus intensities, which then are used to calculate tuning curves for the respective neurons (examples of tuning curves for 3 neighbouring electrodes are shown in Fig. 1). As described above, this approach is only possible in cases where the stimulus onset is known. However, similar LFP events also occur spontaneously without any external stimulus (see Fig. 1a before stimulus onset). These LFP events are usually ignored. Nevertheless, the analysis of spontaneous activity plays a crucial role in several neuroscientific fields such as e.g. the investigation of ongoing phantom perception (e.g. tinnitus).

**Figure 1:**
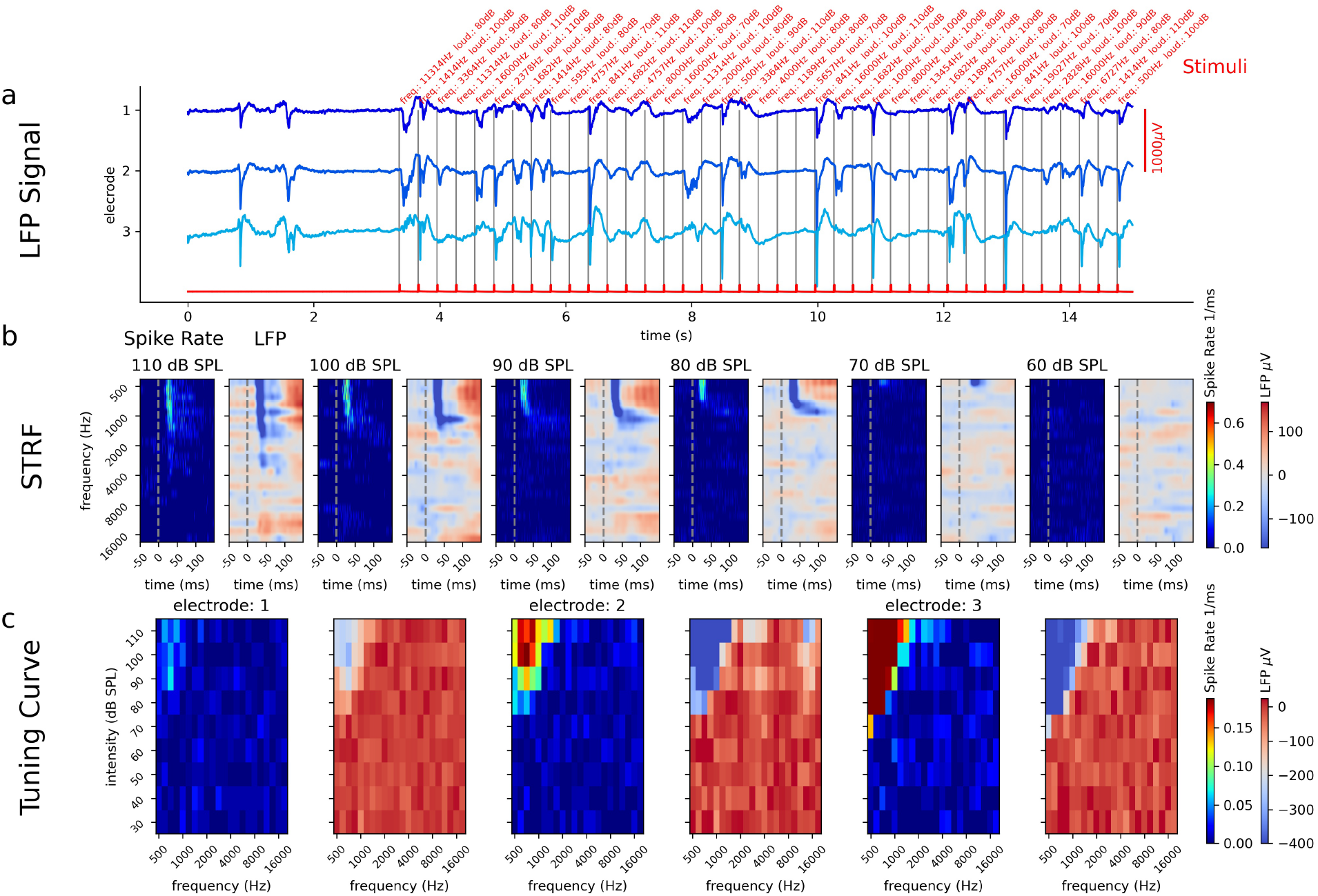
Tuning Curve: a: LFP stream of 3 electrodes (shades of blue) during 50 ms pure tone stimuli (pure tone intensity: 110 dB SPL-70 dB SPL, frequencies: 500 Hz-19027 Hz in half octave steps). The red curve shows the trigger channel (trigger: black vertical lines). b: Spectro-Temporal-Receptive-Fields (STRFs) of position/electrode 3 in terms of spiking activity and field potentials (sound intensities 110 dB SPL-60 dB SPL). c: Tuning-curves calculated from STRFs in b.

Indeed, LFP events also occur spontaneously, without any external stimulus (spontaneous activity measured at three neighbouring positions shown in Fig. 2a). As a first step, these LFP events can be detected e.g. by an algorithm searching for local minima in the ongoing data stream. Additionally, invalid events may be filtered out by defining a threshold and removing events, where the latency between the events is too low. Thus, only the lowest minimum is taken into account (detected LFP events marked by black crosses in Fig. 2a). A huge advantage of the local-minima-search-algorithm is the fact that the detected minimum already defines an exact time-point, which could for example be used to align and average multiple events. Furthermore, the extracted LFP events can be analyzed in terms of shape, size, time of occurrence, etc. (for an example of extracted LFP events see Fig. 2). Thus, the data of the exemplary animal indicates that the inter-event-intervals (time between occurrence of two subsequent events) are distributed in an asymmetric (e.g. log-normal distributed) way, whereas the peak-to-peak amplitudes are more Gaussian-like distributed. Furthermore, the inter-event-intervals show a bi-modal distribution with 2 peaks at different time-points (approx. 1 s and 2 s). Besides these basic measures of the LFP events such as amplitude or inter-event-interval duration, the LFP events contain much more information. For instance, the shape of the events can provide further information on how information is processed in the cortex.

**Figure 2:**
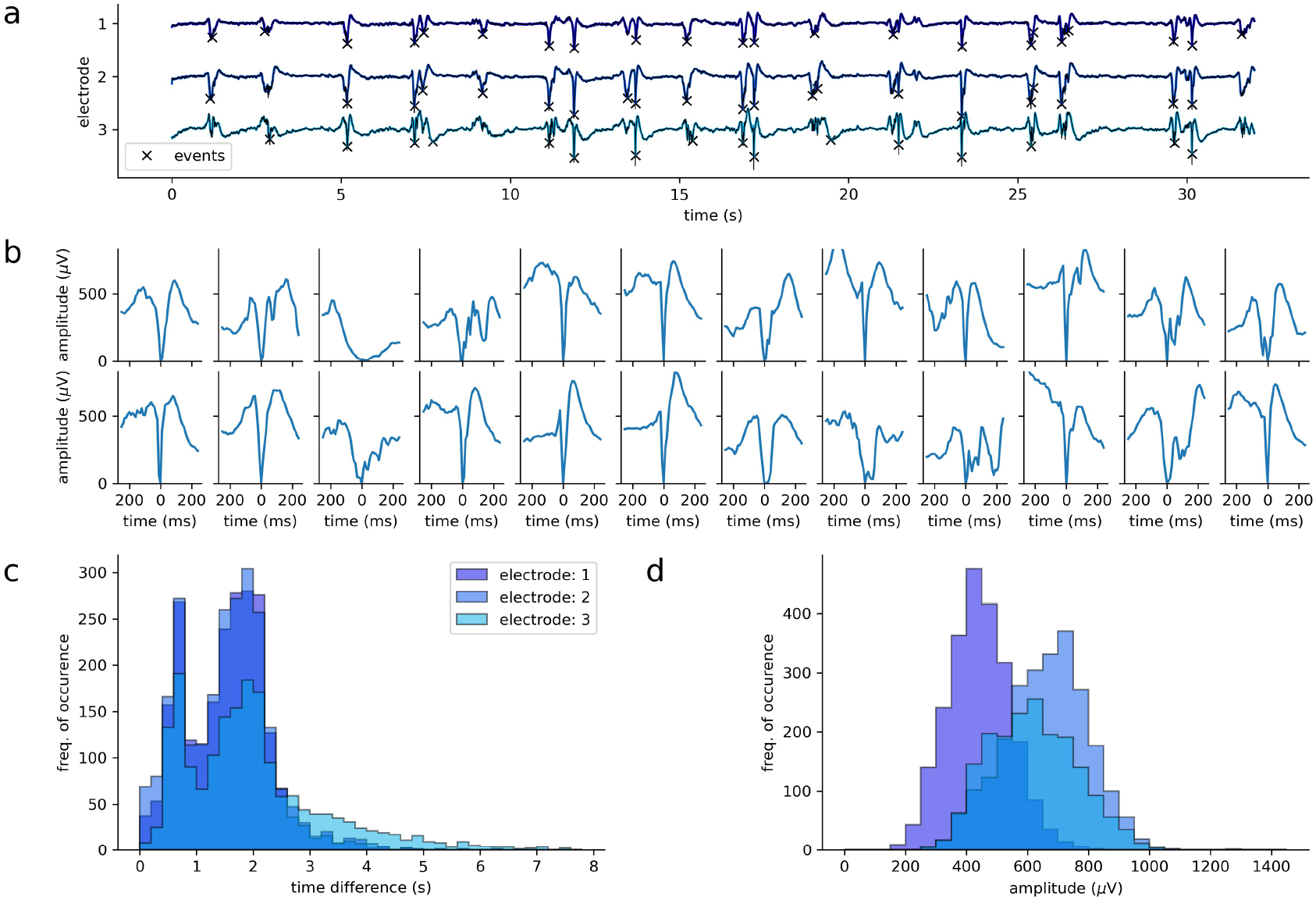
LFP events in spontaneous activity. a: Spontanious activity of 3 electrode channels. Black markers (X) show which events were detected as relevant LFP events. LFP events are detected via local minimum search combined with thresholding; b: Examples (n=24) of detected LFP events; The shape of the events differs. c: Distribution of intervals between detected events; d: Peak-to-peak amplitude distribution of events

### Calculation of embeddings (latent space encodings) of LFP events

In order to extract meaningful information from the LFP events (e.g. the shapes of LFP events) it is necessary to find an efficient representation of these shapes.

Therefore, we calculated so called embeddings by setting up an auto-encoder, i.e. an encoder-decoder-network. This type of artificial neural network has a special network architecture, where the input layer and the output layer are of the same dimension, i.e. consist of equal number of neurons. The network is trained in a self-supervised way, i.e. on reproducing the input in the output layer. Thus, the cost function is the mean-squared error between input and output (schematic drawing of an auto-encoder in Fig. 3a). A further important property of the auto-encoder is the so called bottleneck layer (green layer in Fig. 3a). This means that the intermediate layer is smaller, i.e. has fewer neurons, than the input and output layer. Hence, the auto-encoder has to compress (encode) the input, and subsequently to decompress (decode) the activity in the bottleneck layer in order to reconstruct the input as output. The activation patterns of the bottleneck layer are called latent space embeddings (encodings). The idea behind this processing principle is, that the network has to reduce the dimensionality of the input without loosing relevant information for subsequently reconstructing the input as output again (cyan neurons in 3a). The performance of a given auto-encoder can be assessed by comparing input and output (Fig. 3a). Over-fitting during training the network is prevented by adding dropout layers and testing the trained network) with an unknown test-data set (see Fig. 3b).

**Figure 3:**
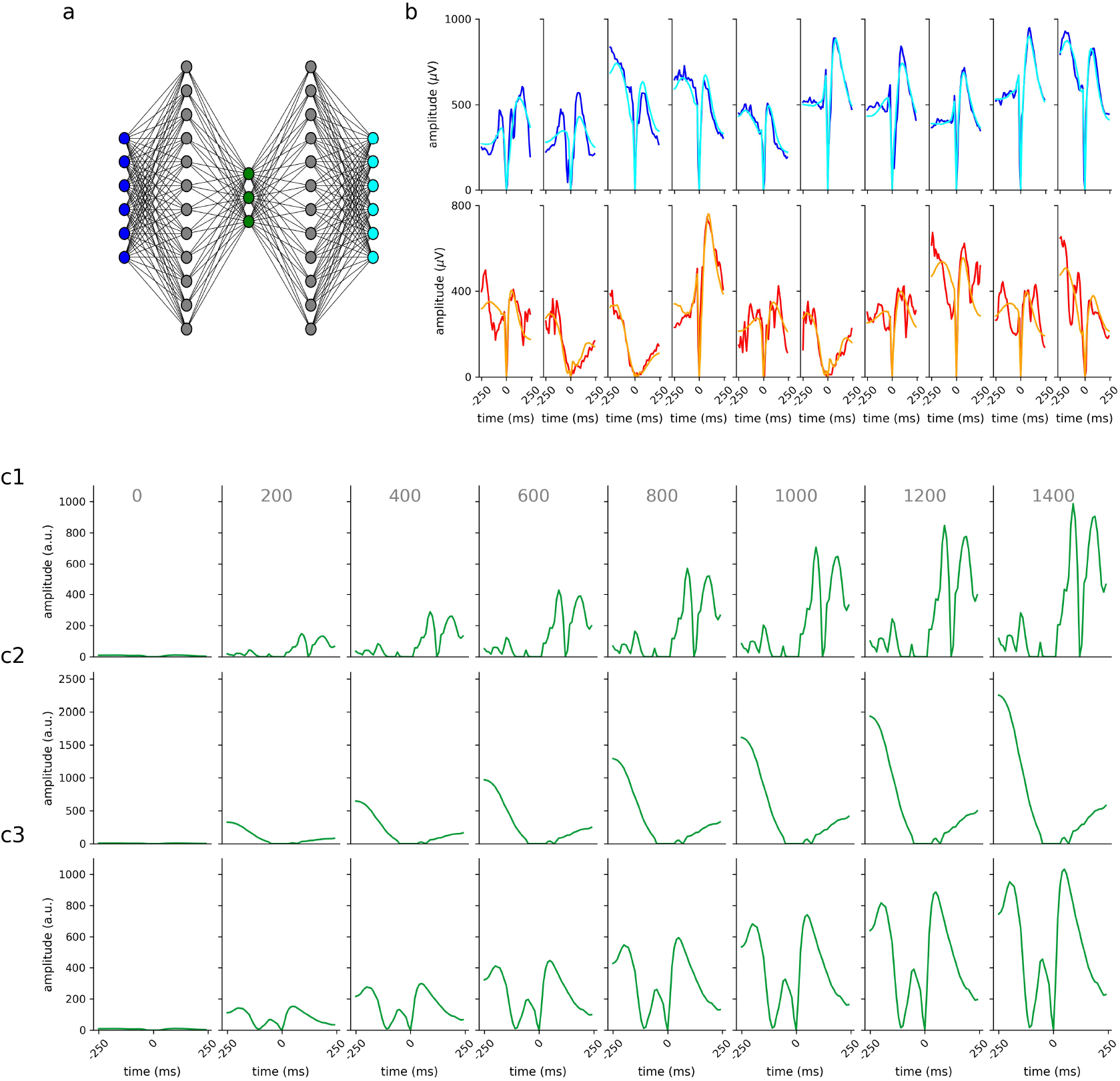
Auto-Encoder for LFP events. a: Scheme of the used auto-encoder for data compression with input layer (dark blue), hidden layers (gray), bottleneck/encoding layer (green), and output layer (cyan). b1: Exemplary input training data (blue) and corresponding outputs/reconstructions (cyan). b2: Exemplary input test data (red) and corresponding outputs/reconstructions (orange). c1-c3: Output of the auto-encoder, for different unity vector activations in the bottleneck/encoding layer, (c1: (x,0,0), c2: (0,x,0), c3: (0,0,x) x∈ {0, 200, 400, 800, 1000, 1200, 1400}). The resulting outputs (c1-c3) represent different degrees of expression of three fundamental complementary event shapes. Any concrete LFP event shape correspond to a weighted superposition of these three prototype shapes.

However, auto-encoders are black boxes and in general it is not known which features of the input space are used to create the lower-dimensional latent space, i.e. what is the meaning of a particular dimension (neuron activity). Therefore, in order to make the encodings more interpretable, we systematically analyzed the emerging latent space embeddings. To this end, we directly activated the neurons in the bottleneck layer one by one without providing any input to the network. In particular, the activation vectors of the bottleneck layer were ((x,0,0), (0,x,0), or (0,0,x) for x∈ {0, 200, 400, 800, 1000, 1200, 1400}). These activation vectors were further propagated through the network to the output layer. The resulting outputs (Fig. 3c1-c3) indicate that each encoding neuron is specialized to certain LFP shapes. For example, neuron 2 seems to encode low-frequency LFP events, whereas neuron 3 is associated with high-amplitude W-shaped signals.

The dimensionality reduction through the auto-encoding can be used to analyze large electrophysiological data sets such as recordings of spontaneous activity for e.g. 1 hour (Fig. 4a-c). Note that, the encodings do neither depend on prior knowledge nor any labeled data. Actually, the auto-encoder only uses statistical features of the input such as the frequency of occurrence of certain LFP events. However, the encodings do not contain the full information of the underlying LFP event shapes, as certain information which is crucial to recosntruct the input is also stored in the decoding part of the neural network itself. Nevertheless, when certain clusters are identified, the shapes of this LFP events can be reconstructed using the decoder (examples of decoded inputs in Fig. 4d-f). The two processing steps: (1) finding certain clusters in the embeddings, and (2) identifying the underlying LFP event shapes, provide efficient data processing on the one hand, minimize the information loss and preserve biological interpretability.

**Figure 4:**
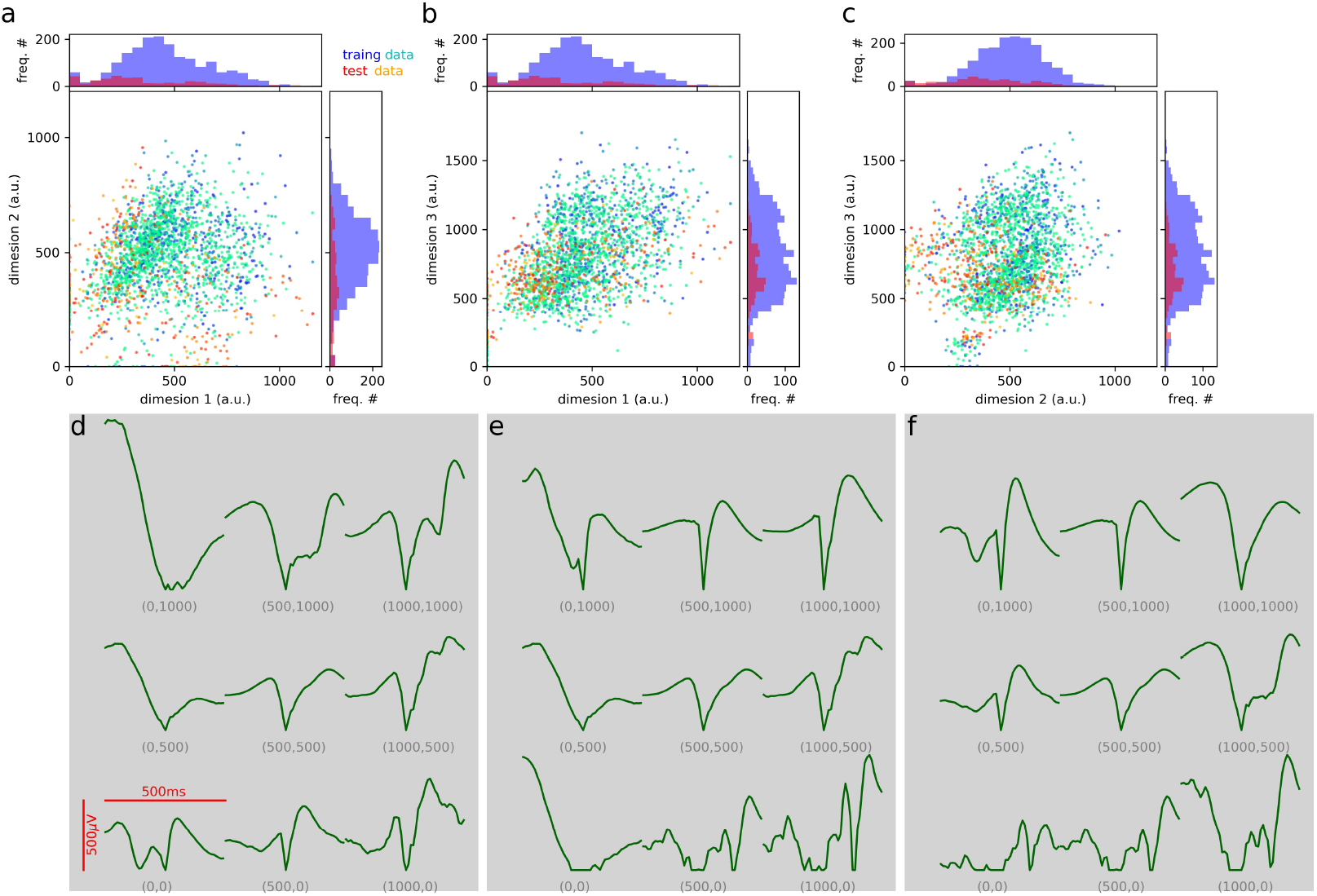
Dimensionality reduction using the auto-encoder. a-c: 2-dimensional projections of the encoded 3-dimensional embeddings of LFP events for the training (blue-cyan) and the test data set (red-orange). Training and test data is spontaneous activity recorded for 1 h or 0.5 h respectively. The time of occurrence of the detected LFP events is color coded (blue to cyan for training data, red to yellow for test data). d-f: Histograms show the distribution of the encoded LFP events. The shapes of the LFP events for different encoding vectors (x,y,z-values ∈ {0, 500, 1000}). Note that, the plots do not show the exact reconstructed curve shapes since they are based on only 2 dimensions, whereas the embedding vector is 3-dimensional. For each column (a/d, b/e, c/f), we have set the respective missing third dimension to a constant medium range value of 500.

As the auto-encoder is exclusively trained on statistical features, the trained network can be applied to any kind of LFP data. Thus, we tested the auto-encoder trained on spontaneous activity to find clusters in evoked activity. Therefore, we used the evoked activity shown in Fig. 1 to check, if the auto-encoder actually helps to extract meaningful information. We analyzed 4 different sound pressure levels (110 dB–80 dB) and 3 frequencies (500 Hz, 1000 Hz, 2000 Hz). We could show that indeed the LFP responses induced by different stimulus frequencies form clusters (Fig. 5a1-d1, for exemplary events and decoded LFP events see a-d 2-4). We quantified the quality of the clustering by calculating the generalized discrimination value (GDV) [50, 51]. We could show that the the GDV is best for a stimulus intensity of 100 dB. Thus, too loud stimuli lead to unspecific activation due to recruiting of auditory nerve fibers. In contrast, too low stimulus intensities do not evoke clear responses. To put it in a nutshell, the auto-encoder trained on a different data set has proven to serve as universal tool to identify clusters of LFP events carrying the information on the stimulus frequency.

**Figure 5:**
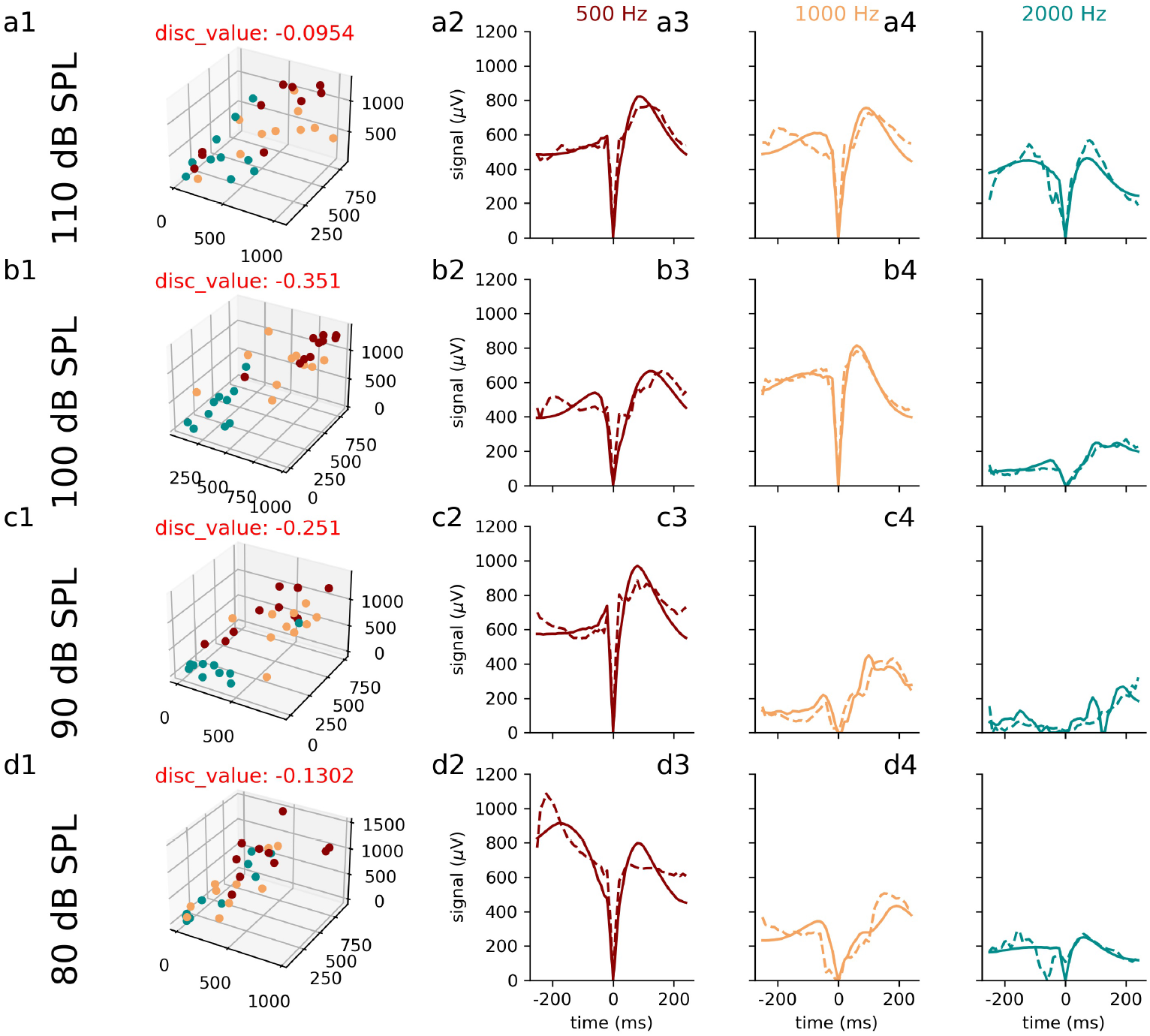
Auto-Encoded LFPs from pure-tone stimulation. Column 1: LFP event shapes in 3-dimensional embedding space. Markers represent encoded LFP responses induced by pure tone stimuli (500 Hz, 1000 Hz, 2000 Hz, best frequency at 500 Hz, compare Fig. 1 electrode 3) for different stimulus intensities, i.e. sound pressure levels (a: 110 dB SPL, b: 100 dB SPL, c: 90 dB SPL, d: 80 dB SPL). The best separability, i.e. lowest generalized discrimination value (GDV, shown in red above plots) of the events can be observed for a optimum stimulus intensity of 100 dB. Louder stimuli cause further recruitment and the specificity to the stimuli is reduced, whereas lower stimulus intensities evoke a decreased signal intensity. Columns 2-4: Examples of input and corresponding reconstructed output shapes for the LFP events shown in a1-d1.

### Estimation of Information Loss Caused by Auto-Encoding using a Classifier Network

However, the dimensionality reduction by auto-encoding (Fig. 6a) obviously causes a loss of information which needs to be quantified. In a previous study, we already introduced a classifier network as a tool to objectively quantify the amount of meaningful information of certain data [**?**]. Training a classifier network (Fig. 6b) requires labeled data. We used a convolutional neural network which was trained on LFP responses induced by four different stimulus frequencies (500 Hz, 1000 Hz, 2000 Hz, 4000 Hz at 100 dB). To quantify the information loss due to auto-encoding/dimensionality reduction with respect to discriminability of different categories of LFP responses, two classifier networks were trained on stimulus frequencies. The original LFP events served as training input for the first classifier, whereas the second classifier was trained on the LFP events which were encoded and decoded by the auto-encoder. Note that, this auto-encoder was trained on data from a different animal (see Fig. 6c). Indeed, the reconstructed input was of sufficient quality (see Fig. 6d).

**Figure 6:**
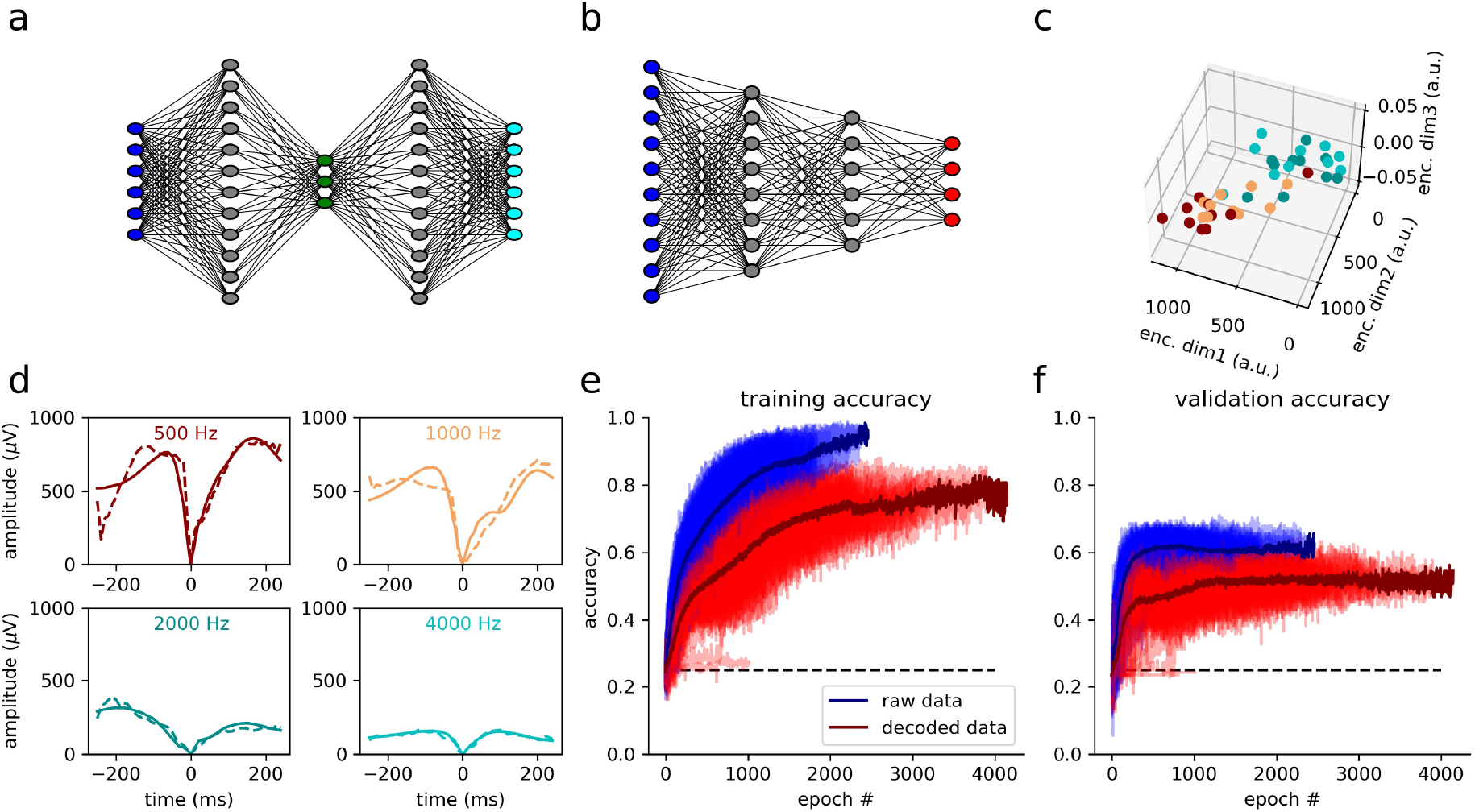
Measurement of the meaningful information of auto-encoded events using a classifier network. a, b: Schematic representation of the two artificial neural networks. Note that, depicted network architectures are sketches, for detailed network parameters see tab. 1. a: Auto-encoder network (compare Figure 3a). b: The classifier network is trained on predicting the frequency (500 Hz, 1000 Hz, 2000 Hz, 4000 Hz) that was used as stimulus and evoked the LFP event shape provided as input to the network c: Reconstructed embeddings of one test data set (10 trials with 4 stimulation frequencies at a stimulation intensity of 100 dB SPL). d: Exemplary LFP responses for different stimulus frequencies (dashed line: measured signal, solid line: reconstructed signal from auto-encoder embeddings). e, f: Training accuracy (e) and validation accuracy (f) of the trained classifier network (dashed line: chance level, dark red/dark blue: average accuracy, red/blue: learning curves for 50 repetitions). The validation accuracy is slightly reduced due to the information loss caused by auto-encoding.

We could show that the classification accuracy (classes correspond to different stimulus frequencies) is slightly reduced for the reconstructed input compared to the unprocessed original input. Nevertheless, there is no large difference between validation accuracies (Fig. 6f). Furthermore, the auto-encoded input data lead to less over-fitting, as the neural network is not trained on specific features or artifacts unique to certain trials (see Fig. 6f). These features get lost during the process of auto-encoding since they are not relevant, i.e. carry no meaningful information, for reconstructing the input from the latent space embedding. Our analysis provides evidence that the information loss due to auto-encoding is acceptable and that the auto-encoder is a valid tool to reduce the dimensionality of the data. Up to now, the auto-encoding procedure was only applied to single trials of single channel data. In a next step, we show that the encoded data can be used to draw conclusions about information processing in the brain.

### The Relationship of Spontaneous and Evoked LFP events

We show that the embeddings of the LFP events evoked by pure tones form a subspace of a larger state space, which is spanned by the spontaneous LFP event embeddings (see Fig. 7). Thus, during spontaneous activity the brain samples from the realm of possible stimulus evoked event shapes. This phenomenon was already described by Luczak and colleagues in 2009 [52] in the context of population coding. In their study, they analyzed multi-channel spiking activity of several neurons recorded with multi-electrode arrays. However, in our study, we confirm their findings by analyzing cortical activity patterns through the shape of single-channel LFP events.

**Figure 7:**
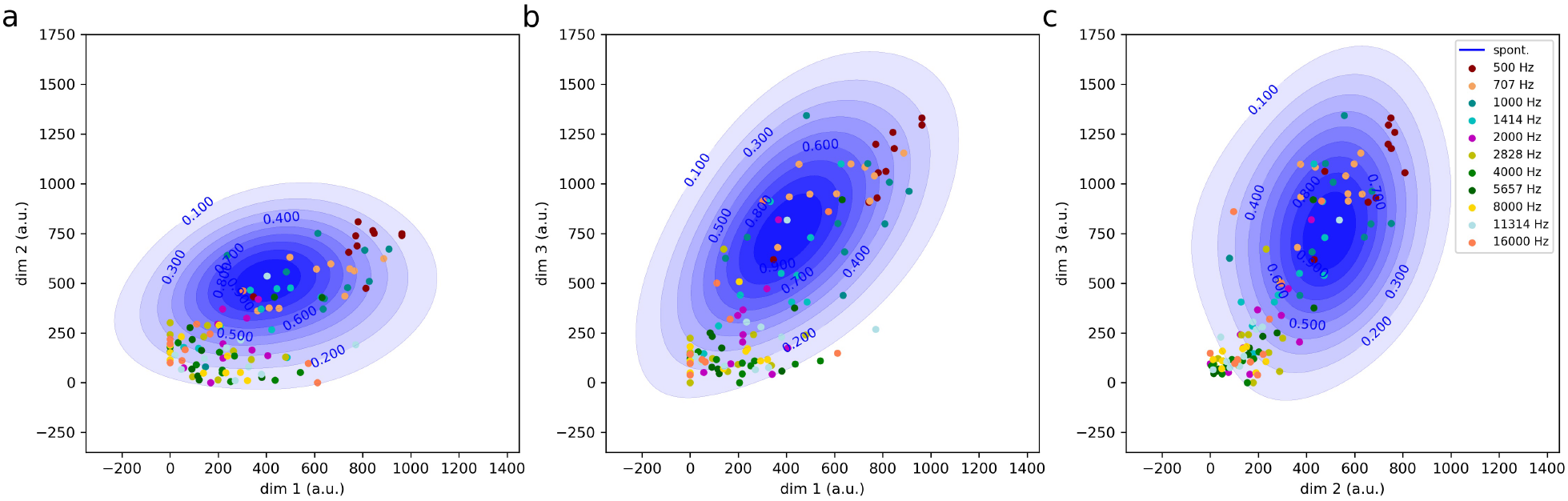
Spontaneous LFP events outline the realm of possible evoked LFP events. Shown are three 2D projections of the 3D embedding space (a: dim 1 and dim 2, b: dim 1 and dim 3, c: dim 2 and dim 3). Markers represent auto-encoder embeddings of LFP events evoked by auditory stimuli of different frequencies (500 Hz-16 kHz, half octave steps, 100 dB SPL, 10 repetitions for each frequency). Spontaneous activity is shown as contour plots (blue) representing the kernel density estimation of spontaneous LFP events.

So far, we restricted our analyses on LFP events from a single recording channel. In the following, we show that we can even derive information about the temporal dynamics of information flow from LFP events by taking into account two spatially separated recording channels.

### Auto-encoding, Clustering and Information Flow

Therefore, we detected spontaneously occurring LFP events in a certain reference channel and analyzed the activity in a spatially separated, neighboring channel (example for events in reference channel in Fig. 8a blue curve and neighbouring channel green).

**Figure 8:**
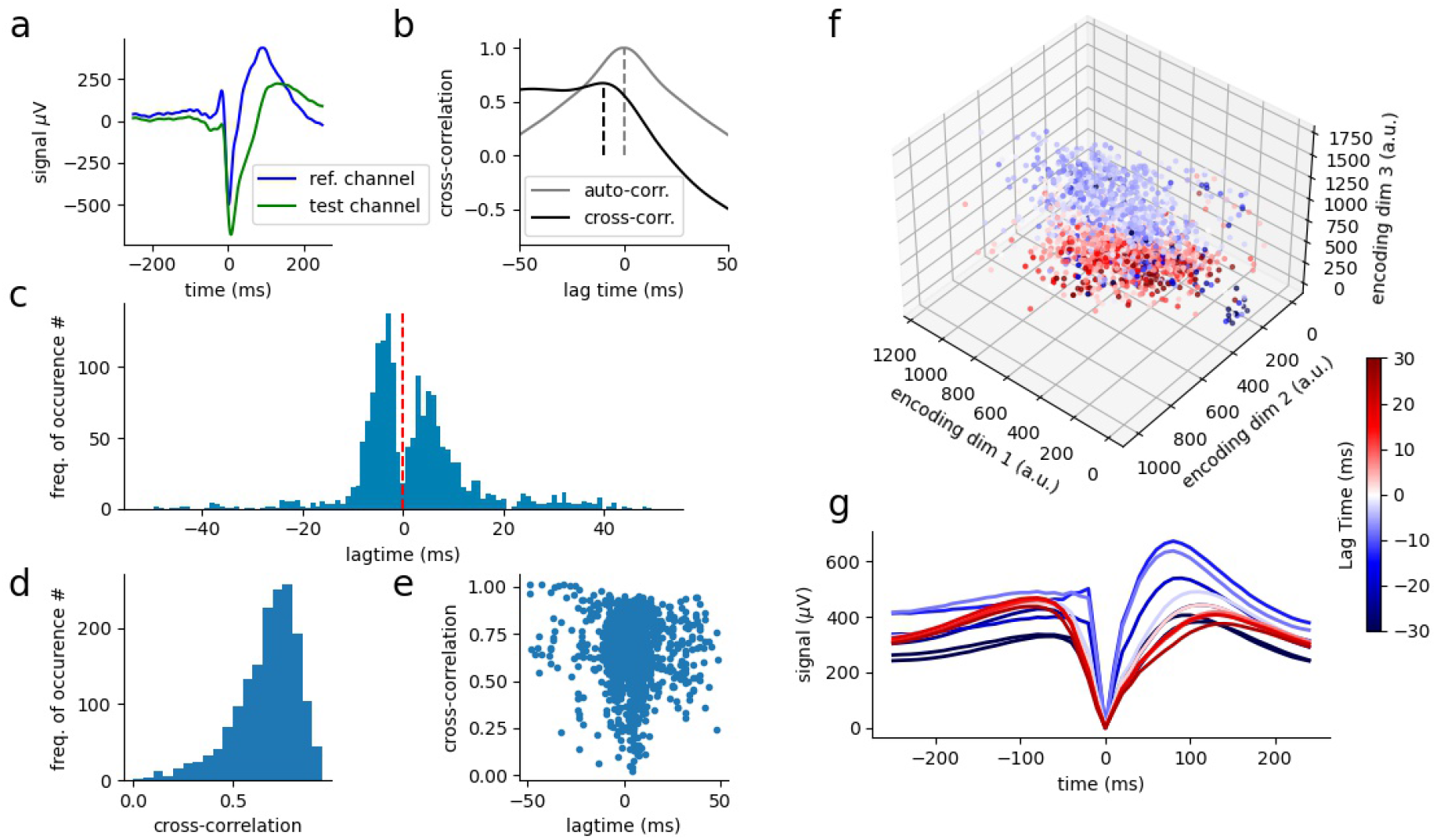
Event-wise cross-correlation of spontaneous LFP events. a: LFP events in reference (blue) and in a neighbouring channel (green). b: Auto-correlation of reference-channel (gray) and cross-correlation between channels (black). c: Histogram of detected lag-times. d: Histogram of maximum cross-correlation coefficients. e: Scatter plot of lag-times and corresponding cross-correlation coefficients. f: Encoding of all events of reference channel. Colors represent different lag times between reference channel and neighbouring channel. g: Reconstructed event shapes from embeddings shown in f. Again, colors represent different lag-times as shown in f (blue: negative lag-times, red: positive lag-times). The sign of the lag-time indicates direction of information flow. Thus, blue represents input from thalamus and red represents input from other cortical areas. The LFP event shape embeddings cluster according to lag-times. Sharp asymmetric curves are related to thalamic input (blue curves g).

The the sign of the time difference between corresponding LFP events in two different channels indicates the direction of the information flow between the two recording sites. To quantify the time difference (latency) between corresponding events, the event-wise cross-correlation function between the two channels of interested was calculated (see Fig. 8b black curve). For each pair of events in the two channels, the lag-time which leads to the maximum cross-correlation coefficient corresponds to the respective time difference between the occurrence of the same event at the two different channels. It turns out that there is no general lag-time/latency value that fits to all pairs of events. Instead, spontaneous activity is characterized by a distribution of latencies as shown in Fig. 8c. This means that simply calculating the cross-correlation function over the complete LFP time series or spike trains of two recording channels as is frequently done as an ojective function of the synchrony between two channels is insufficient. For instance, an increased synchrony could either be caused by a more balanced distribution of negative and positive time delays (lag-times) between corresponding LFP events, or by smaller absolute values of the time delays. This information gets lost by simply calculating cross-correlation functions for the entire time series of both channels. Furthermore, it turns out that the resulting cross-correlation coefficients are also broadly distributed between 0 and 1 (Fig. 8d), whereas there seems to be no clear dependence between resulting lag-time and maximum correlation coefficient. In Figure 8e, a scatter plot of all pairs of cross-correlation coefficients and corresponding lag-times is shown. This indicates that the value of the cross-correlation coefficient is no reliable measure for the synchrony between two channels.

We took advantage of the previously calculated latent space embeddings of LFP event shapes to assess, if the time delays of LFP events between two channels correspond to the shape of the respective LFP event. Remarkably, the event shape embeddings, and consequently the event shapes, cluster according to the direction of information flow, i.e. the sign of the lag-time between two channels (Fig. 8f). Finally, we decoded the embeddings again to visualize the corresponding LFP event shapes of the channel of interrest. We find that LFP event shapes characterized by larger amplitudes correspond to negative (blue) time delays, whereas broader shapes with smaller amplitudes correspond to positive (red) time delays, and hence indicate information flow in the opposite direction. Thus, it is possible to estimate the direction of information flow in the cortex by analyzing only a single recording channel. These findings indicate that it might be possible to assess changes in information flow direction, when the system is distorted e.g. by damages along the sensory pathway such as hearing loss caused by a noise trauma.

Indeed, noise trauma leads to changes of the information flow as seen in Fig. 9. Depending on the electrode position, the embedding clusters changed when the silence condition (9a,b 1-2) is replaced by a 115 dB, 2 kHz auditory pure-tone trauma (9 a,b 3-4).

**Figure 9:**
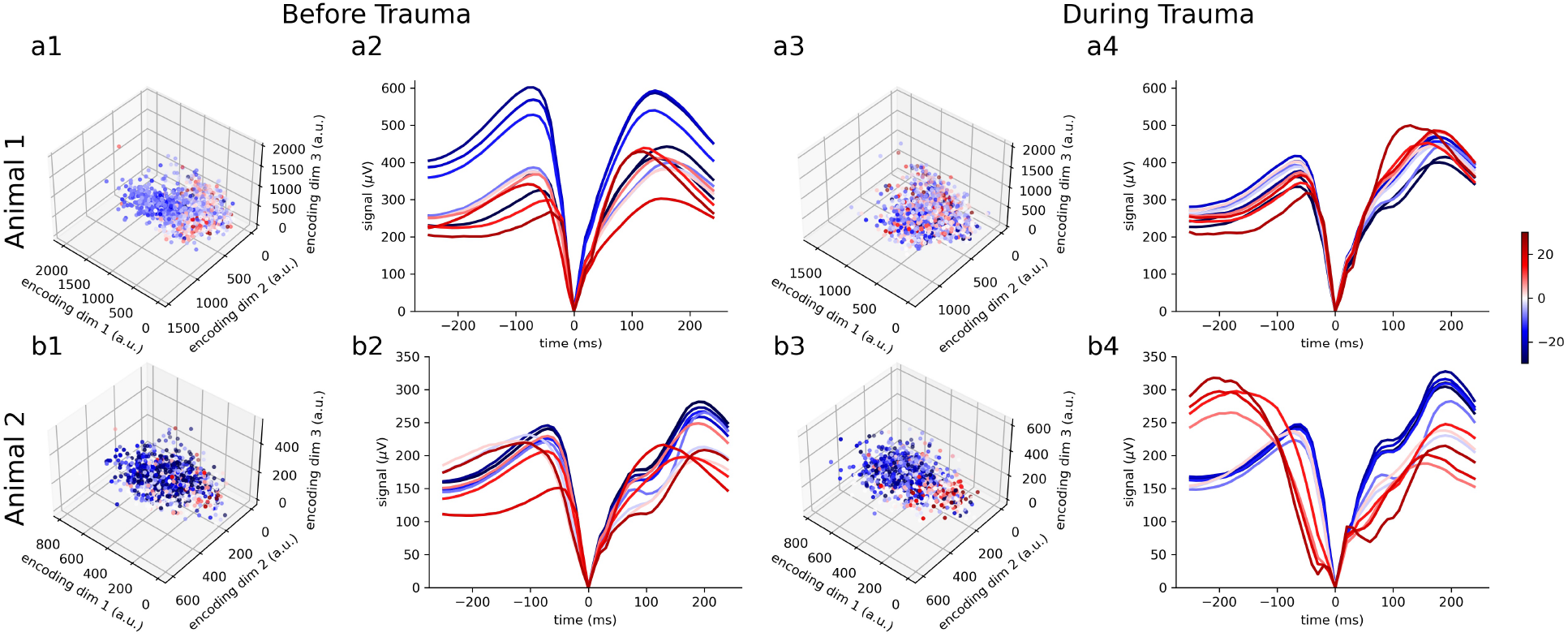
Event-wise cross-correlation of spontaneous LFP events before and during noise trauma. . a1: LFP event shape embeddings from an exemplary animal before noise trauma. a2: Reconstructed event shapes of embeddings from a1. a3-a4: Embeddings (a3) and corresponding reconstructions (a4) of LFP event shapes of the same animal shown in a1 and a2 but during application of an auditory noise trauma (2 kHz pure tone, 115 dB SPL, 60 min); b1-b4: Same as a1-a4 for a second animal.

### Application of the method to human intra-cranial EEG (iEEG) data

The fact that the information flow can be determined by analyzing single channel LFP event shapes could be interesting for analyzing human intra-cranial EEG (iEEG) data. We applied the auto-encoding procedure on iEEG data recorded in the the auditory cortex of a human epilepsy patient. Indeed it is possible to use the local-minimum search algorithm on the recorded data to identify LFP events (see 10a,b, events are marked by x). Besides spontaneous activity also evoked activity was recorded: currents between 1 mA and 15 mA were applied to certain channels of the recording device. We distinguish between stimulation of channels, which are not our recording channels (see red markers in Fig. 10a) and currents which cause artifacts (green markers Fig. 10a). Note that, in principle, events labeled as spontaneous events might nevertheless be evoked events since we cannot fully exclude that there were no sounds or other auditory stimuli in the room during recording. Thus, for this analysis evoked events are defined as events which are evoked by an intra-cranially applied stimulation current. We find that auto-encoding is again a valuable technique to extract meaningful data from the iEEG data.

**Figure 10:**
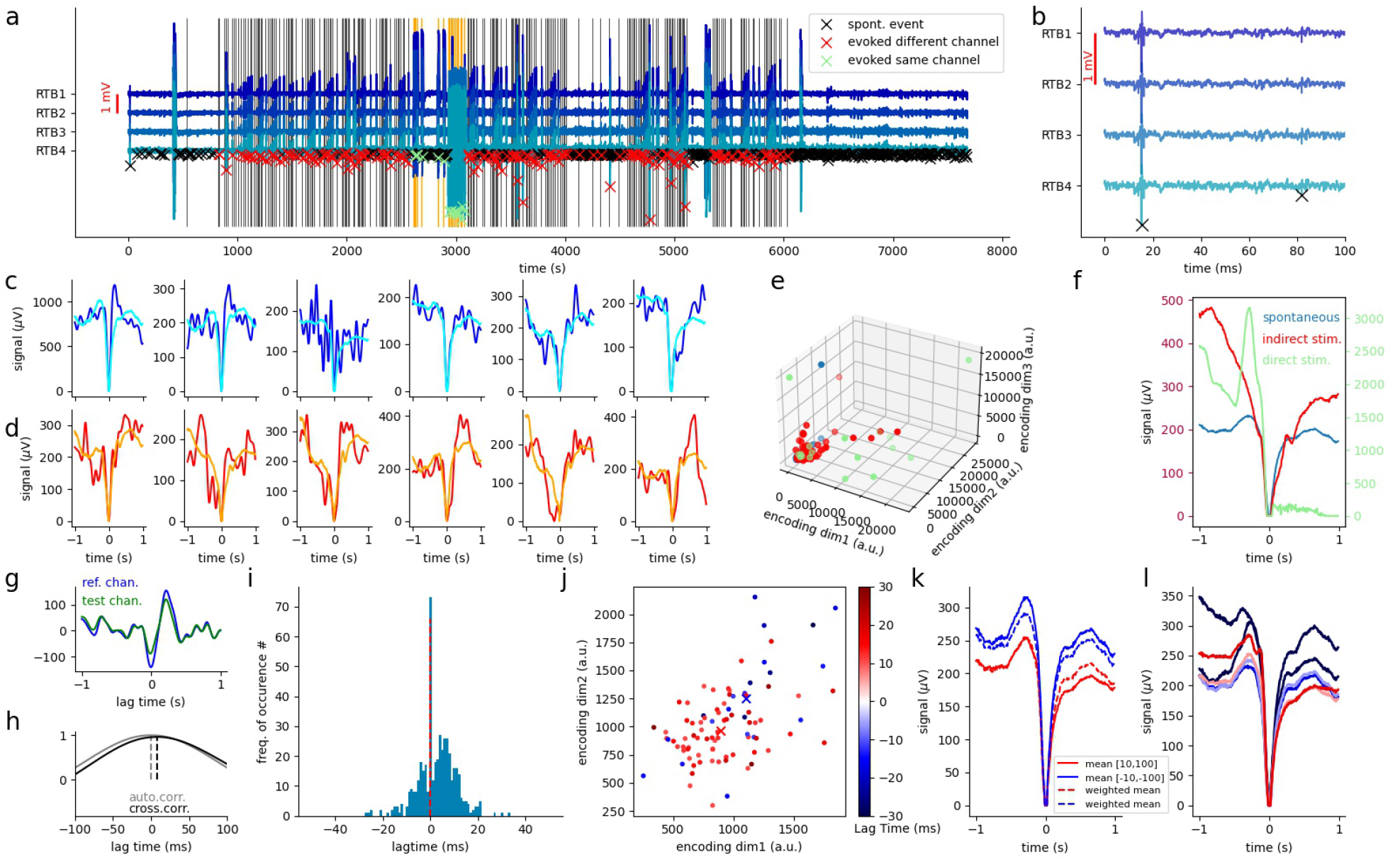
Application of auto-encoding approach on human iEEG data. a: Intracranial recording (iEEG) in the auditory cortex of a human epilepsy patient (shown are four channels: RTB1-4). Markers (black, green, red) indicate LFP events identified by the local minimum search algorithm (see Methods). Red markers: Events evoked by stimulation currents induced in electrodes different from RTB1-4. Green markers:Events which were directly induced by current induction into the electrodes RTB1-4. Black markers: Spontaneous or auditory evoked events. b: Temporal zoom of time series shown in a. c, d: Examples of spontaneous LFP event shapes and corresponding reconstructions from auto-encoder embeddings of the training data set (blue: input, i.e. original recorded LFP events, cyan: reconstructed LFP events from embeddings) and the test data set (red: input LFP events, orange: reconstructions from embeddings). e: Embeddings of LFP events (blue: spontaneous, green: evoked by current in channels RTB1-4, red: evoked by current in different channel). f: Reconstructions calculated from median embeddings (from e) of the three different conditions. Note that, the reconstructions represent prototypical LFP shapes for the different conditions. g, h: Examplary LFP events of channel RTB4 (green, reference channel) and channel RTB3 (blue, test channel) and corresponding cross correlation (b, black curve). The gray curve shows the auto-correlation of the reference channel. i: Distribution of the lag-times between the two channels (position of black peak). j: Embeddings of all spontaneous LFP events with lag-times smaller than −10 ms (shades of blue) or larger than 10 ms (shades of red). Markers (x) represent the average embeddings (blue cross: average of LFP events with latency smaller than −10 ms, red: > 10 ms). k: Reconstructions correspodning to the mean embedding coordinate values shown in j (solid line: mean of LFP events with lag-times *in*[−100, −10] ∪ [10, 100], dashed line: weighted mean over all events). l: Systematic reconstructions for different lag times.

First, we demonstrate that the auto-encoder network produces valid reconstructions as output compared to the input data (see Fig. 10c,d). Furthermore, evoked events lead to different LFP shapes than spontaneous LFP events. Even though, no distinct clusters in the embedding space could be observed, the median coordinate values of the three different conditions lead to three different reconstructed prototypical LFP event shapes (see Fig. 10f).

The inter-channel cross-correlation analysis (Fig. 10g,h) indicates that LFP event shape are correlated with lag-times between the channels. Thus, we find again a bi-modal lag-time distribution (Fig. 10i). Furthermore, a correlation between LFP event shape and lag-time can be observed, at least for lag-times with an absolute value larger than 10 ms, (Fig. 10j). Thus, the reconstruction of the average (prototypical) embeddings for lag-times larger than 10 ms and smaller than −10 ms indicate that at least the amplitude of the LFP events with negative lag times are increased (see also corresponding animal data in Fig. 8g and 9a2, b4).

## Discussion

In the present study, we developed an analysis pipeline to identify, extract and characterize events from ongoing recordings of local field potentials (LFP). We applied a local minimum search algorithm in combination with a thresholding procedure to identify significant LFP events. In a next step, the dimensionality of the LFP event shapes is reduced using an auto-encoder network. In its bottleneck layer, the auto-encoder provides a low-dimensional representation (embedding) of the input data, which conserves relevant information to reconstruct the original shapes again as output. These embeddings are used to visualize the data and to identify potential stimulus-related clusters. Note that, the clusters result from the properties of the electrophysiological recordings and are not due to any pre-defined labels. The embeddings were further used to show that the shape of the LFP events is correlated with the direction of the information flow between different recording sites/channels (see Fig. 8). In our example, sharp high-amplitude LFP events indicate that the source of the LFP event is located at the respective channel, whereas broad low-amplitude shapes indicate that the source is at the other channel. This means that LFP event shapes can be used to identify the location of sub-cortical input. However, auto-encoder based dimensionality reduction leads to worse interpretability of the underlying LFP event shapes, as the low-dimensional representations are highly abstract and auto-encoders, as deep learning in general, suffer from the so called black box problem [53]. Therefore, we calculated prototypical embeddings by averaging over all embeddings belonging to a certain stimulus conditions. Subsequently, we reconstructed the corresponding typical LFP shapes. These prototypical LFP event shapes could be used to make further assessments on cortical information processing. For instance, we obtained the remarkable result that during spontaneous activity without any stimulus, LFP event shapes ae sampled from the realm of possible stimulus evoked LFP event shapes. A phenomenon that has so far only been demonstrated in the context of multi-channel spike train population coding [52].

In summary, the benefit of using auto-encoders for processing LFP data can be divided into three major points. First, the dimensionality reduction allows for visualizing highly complex data sets. The fact that the embeddings can be reverse-engineered to prototypical LFP event shapes increases interpretability. Furthermore, there is another significant advantage of this procedure. The auto-encoder can be used to de-noise the data: as the auto-encoders are exclusively trained on statistical features of the LFP events, unique artifacts do not play a significant role. Thus, the encoding-decoding procedure can be used for artifact suppression (for some existing approaches see [32, 54–56]). A further central finding of the study is, that lag-times of LFP events measured between two channels are broadly distributed and this distribution is bi-modal, i.e. with two maxima for positive and negative lag-times, respectively (see Fig. 8c and Fig. 10i). Our data indicates that the different LFP event shapes correspond to different information processing mechanisms. Since the analysis of neural synchrony is an important target to investigate auditory processing in the cortex, and is of particular meaning for different phantom perceptions such as tinnitus theories [57, 58], this finding could be a starting point for further investigations. Indeed in most studies, synchrony between LFP streams from different channels is quantified by calculating the cross-correlation function between the entire signal streams, analyzing lag-times and the area under the cross-correlation curve [59]. However, our findings that lag-times correspond to different LFP event shapes and that lag-time histograms show two maxima indicate that standard synchrony analyses fail to provide the full picture. Applying the here presented novel approach to further investigate the functional plasticity after hearing loss, which is hypothesized to be the cause of tinnitus [60–66], might lead to a deeper understanding of the underlying processes.

However, analyzing LFP-events using auto-encoder networks has also some drawbacks, that have to be taken into account when interpreting the results. As we use the decoder part of the auto-encoder to generate reconstructions from average embeddings, we create novel LFP shapes and thus the decoder part of the network works as a generative model [67]. Overfitting is an important problem for all machine learning applications [68,69] and especially for generative models. When the auto-encoder network has too many trainable parameters negative side effects can occur. For instance, the network could store the whole information about different LFP shapes within the large weight matrix [70]. Therefore, the weight matrix would serve as some kind of list or look-up table for different LFP shapes, and the embedding layer would just learn random labels (indices) for the content of this list. This unwanted case would cause the effect that the embeddings alone do not contain any useful information about the LFP shape, because it would be possible to train further auto-encoder with arbitrary permutations of the embeddings, which perform equally good. However, in order to allow an interpretation of clusters in embedding space, neighbourhood relations between different LFP shapes should be conserved in the embedding layer [71]. Thus, in our study we used shallow-networks to reduce the number of parameters, and added drop-out layers to prevent the neural network from overfitting. We could show that the encoder does not simply add labels to the different LFP-shapes, because the self-organized emerging clusters actually correspond to different stimulus conditions (see Fig. 5). As the neural network was not trained on these LFP event shapes yet from another data set and the clusters automatically emerge we could show that embeddings are not just random representations of the different LFP event shapes. Nevertheless, in a follow-up study, as a more sophisticated approach, variational auto-encoders [72] or transformer networks [73] could be used instead, which are further optimized to lead to better encondings and thus to more interpretable decodings [74, 75].

Summing up, following the trend of integrating artificial intelligence and neuroscience [76–83], machine learning provides valuable tools to extract information from electrophysiological data [84–86]. As described above in most studies the data is averaged over many measurement trials to increase the signal to noise-ratio. However, the frequently performed averaging procedure erases any correlates of information processing taking place during recording of the ongoing continuous signal stream. Potentially, it is possible to translate that stream of voltage fluctuations into signs, which could be interpreted by humans using approaches from deep learning based natural language processing such as machine translation, or even from *animal linguistics*, where e.g. killer whale calls are identified, segmented, extracted and classified from ongoing continuous sound streams according to recurring feature patterns [87–90]. By that, we are convinced that our approach might further push the progress in neuroscience in order to extract meaningful information from continuous electrophysiological data streams.

## Author contributions

AS and PK conceived the study. PK supervised the study. AS, CB and JR performed the electrophysiological recordings in rodents. HH and CR provided human data. AS and RG wrote the evaluation software. AS, RG, AM, CM, HH and PK wrote the manuscript. All authors approved the final version of the manuscript.

## Acknowledgments

This work was funded by the Deutsche Forschungsgemeinschaft (DFG, German Research Foundation): grant KR 5148/2-1 (project number 436456810) to PK, and grant SCHI 1482/3-1 (project number 451810794) to AS. Furthermore, the research leading to these results has received funding from the European Research Council (ERC) under the European Union’s Horizon 2020 research and innovation programme (ERC Grant No. 810316 to AM).

